# Predator complementarity dampens variability of phytoplankton biomass in a diversity-stability trophic cascade

**DOI:** 10.1101/851642

**Authors:** Chase J. Rakowski, Caroline E. Farrior, Schonna R. Manning, Mathew A. Leibold

## Abstract

Trophic cascades – indirect effects of predators that propagate down through food webs – have been extensively documented. It has also been shown that predator diversity can mediate these trophic cascades, and separately, that herbivore biomass can influence the stability of primary producers. However, whether predator diversity can cause cascading effects on the stability of lower trophic levels has not yet been studied. We conducted a laboratory microcosm experiment and a field mesocosm experiment manipulating the presence and coexistence of two heteropteran predators and measuring their effects on zooplankton herbivores and phytoplankton basal resources. We predicted that if the predators partitioned their zooplankton prey, for example by size, then co-presence of the predators would reduce zooplankton prey mass and lead to 1) increased average values and 2) decreased temporal variability of phytoplankton basal resources. We present evidence that the predators partitioned their zooplankton prey, leading to a synergistic suppression of zooplankton; and that in turn, this suppression of zooplankton reduced the variability of phytoplankton biomass. However, mean phytoplankton biomass was unaffected. Our results demonstrate that predator diversity may indirectly stabilize basal resource biomass via a “diversity-stability trophic cascade,” seemingly dependent on predator complementarity, but independent of a classic trophic cascade in which average biomass is altered. Therefore predator diversity, especially if correlated with diversity of prey use, could play a role in regulating ecosystem stability. Furthermore, this link between predator diversity and producer stability has implications for potential biological control methods for improving the reliability of crop yields.

## INTRODUCTION

A substantial body of work, generally motivated by global biodiversity loss, indicates that enhanced biodiversity of ecosystems often stabilizes community biomass (Jiang and Pu 2009, Loreau and de Mazancourt 2013, Gross et al. 2014). Most of these studies measured or modeled the effects of primary producer diversity on the variability of primary producer biomass (e.g. Hector et al. 2010, Loreau and de Mazancourt 2013, Gross et al. 2014). However, predators face greater extinction threats than do lower trophic levels, suggesting that predator diversity is more relevant to global biodiversity change than is primary producer diversity (Purvis et al. 2000). Furthermore, biodiversity often alters energy flow through food webs, and so manipulating diversity and measuring its effects within a single trophic level can give an incomplete picture of how biodiversity influences ecosystem functioning (Hines et al. 2015, Seabloom et al. 2017). While some literature addresses how predator diversity affects the average biomass of various other trophic groups in food webs (reviewed in Schmitz 2007), little is yet known about the influence of predator diversity on ecosystem stability. In particular, the existence of a link between predator diversity and the stability of non-adjacent lower trophic levels has not (to our knowledge) been tested. Yet, such a link would have critical implications for both the maintenance of stable natural ecosystems as predator diversity changes, and also for biological control to potentially stabilize crop yields.

Existing research relating predator diversity to lower trophic level functioning has shown that higher predator (or natural enemy) diversity generally leads to lower mean herbivore densities and higher mean plant biomass, as long as the predators exhibit complementarity in their feeding niches (Straub et al. 2008, Greenop et al. 2018). This research was mostly performed in the context of biological pest control, but relies on general ecological principles. Therefore, diversity of functional traits among predators related to prey use, such as body size, may generally play a key role in mediating trophic cascade strength (Straub et al. 2008). But besides average primary producer biomass, the temporal variability of primary producer biomass could also be affected by functional predator diversity in a predictable way. While herbivory generally decreases primary producer biomass, it has also been shown to increase the variability of primary producer biomass (Thébault and Loreau 2005, Downing et al. 2014), perhaps resulting from predator-prey cycling between the herbivores and producers (Volterra 1926); though note that the opposite effect has also been found (Post 2013). Thus, if functional predator diversity reduces herbivore biomass, it may also indirectly reduce the variability of primary producer biomass, producing a “diversity-stability trophic cascade” (Table 1). Such an effect would stem from two coupled relationships: 1) a negative relationship between predator diversity and herbivore (prey) biomass, likely due to prey use complementarity, and 2) a positive relationship between herbivore biomass and producer variability.

**Table 1.**
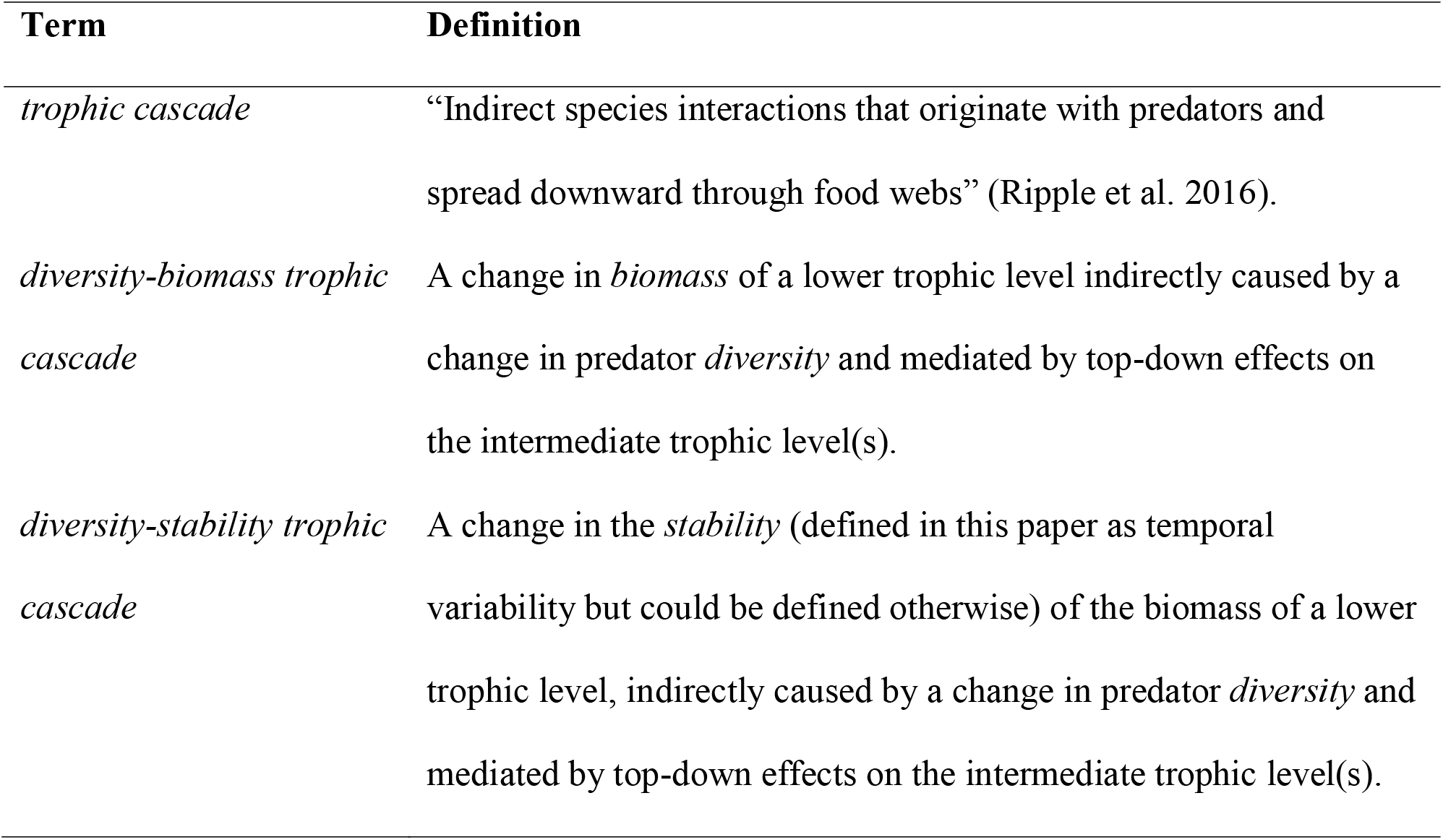
Trophic cascade terminology used in this paper.

| Term | Definition |
| --- | --- |
| *trophic cascade* | “Indirect species interactions that originate with predators and spread downward through food webs” (Ripple et al. 2016). |
| *diversity-biomass trophic cascade* | A change in *biomass* of a lower trophic level indirectly caused by a change in predator *diversity* and mediated by top-down effects on the intermediate trophic level(s). |
| *diversity-stability trophic cascade* | A change in the *stability* (defined in this paper as temporal variability but could be defined otherwise) of the biomass of a lower trophic level, indirectly caused by a change in predator *diversity* and mediated by top-down effects on the intermediate trophic level(s). |

Here we report the results of a laboratory microcosm experiment and a field mesocosm experiment to test for the existence of diversity-stability trophic cascades. After testing for prey use complementarity by two predator species in the laboratory, we manipulate the presence of the same predators in the field (no predators, each predator alone, and both predators) and measure the resulting biomass of plankton groups as well as the variability of phytoplankton biomass. We use the heteropterans *Notonecta* and *Neoplea* as the predators due to their substantial difference in body size and consequent likelihood of partitioning prey resources. Based on previous work (Murdoch et al. 1984) we predicted that *Notonecta* would mostly consume *Daphnia*, and we predicted that *Neoplea* would consume smaller zooplankton based on its smaller body size. We hypothesized that if *Notonecta* and *Neoplea* partition zooplankton prey, then the combination of these predators would suppress zooplankton more than either predator alone, leading to two indirect effects: 1) increased mean phytoplankton biomass (i.e., a diversity-biomass trophic cascade, Table 1b), and 2) reduced variability of phytoplankton biomass (i.e., a diversity-stability trophic cascade, Table 1c). We present evidence that these predators indeed partitioned zooplankton prey, leading to a greater suppression of zooplankton when the predators were together than predicted from their individual effects; and that this suppression of herbivores indirectly stabilized, but did not enhance, phytoplankton biomass. In order words, we detected a diversity-stability trophic cascade despite the absence of a diversity-biomass trophic cascade.

## METHODS

### Focal predators

We used two insect species in the suborder Heteroptera as the predators in both the field and laboratory experiments: the notonectid *Notonecta undulata* and the pleid *Neoplea striola*. Hereafter, we use the genus names (*Notonecta* and *Neoplea*) to refer to these two focal predator species. Both are mobile generalist predators that are widespread across North America. However, they differ starkly in size: *Notonecta* adults measure ∼11-13 mm and *Neoplea* adults measure ∼1.5 mm in length. Studies have shown that while members of the genus *Notonecta* can take prey as small as microscopic rotifers, they strongly reduce large prey such as *Daphnia* and mosquito larvae (Leon 1998, Murdoch et al. 1984, Hampton and Gilbert 2001). The genus *Neoplea* has been less studied; they have been documented to attack invertebrates ranging in size from rotifers to *Daphnia* (Hampton and Gilbert 2001, Gittelman 1977), but we predicted they would prefer smaller prey than *Notonecta*.

### Organism collection

We allowed communities of phytoplankton to assemble naturally in six outdoor tanks at the University of Texas’ Brackenridge Field Laboratory, Austin, TX for ∼six months. We then mixed a common inoculum from these tanks. The phytoplankton community became dominated by green algae (Chlorophyta), ranging in size from green picoplankton ∼1 µm in diameter to *Oocystis* with mother cell walls up to ∼25 µm in diameter and dominated by a few morphospecies, especially *Selenastrum* and *Oocystis* (Appendix S1: Table S1). We collected an array of zooplankton taxa from small water bodies nearby, including many rotifer species, *Spirostomum, Arctodiaptomus dorsalis, Mesocyclops edax*, and *Scapholeberis kingi*, and we ordered *Daphnia magna* from Sachs Systems Aquaculture (St. Augustine, FL) to extend the size range of the zooplankton to larger-bodied individuals (Appendix S1: Table S2). We similarly mixed the zooplankton taxa together into a common inoculum. The predators *Notonecta* and *Neoplea* were also collected from small water bodies in Austin, TX.

### Laboratory experiment

To test whether *Notonecta* and *Neoplea* partitioned zooplankton prey resources, we conducted a short time scale (5-day) experiment in the laboratory. Five days represents just under one generation for the dominant zooplankton with the shortest generation times, *Daphnia magna* and *Scapholeberis kingi*. Therefore we anticipated that five days would provide enough time for the predators to reduce zooplankton populations but would not provide enough time for the zooplankton populations to significantly recover, allowing us to better estimate the effects of the predators on mortality of different zooplankton species while minimizing the influence of zooplankton fecundity and plasticity. First, we combined portions of the phytoplankton and zooplankton inocula into a common mixture. Then we placed ten *Notonecta* adults individually in microcosms with 1.5 L of the plankton mixture, and likewise placed ten *Neoplea* adults individually in microcosms with 100 mL of the plankton mixture. This 15:1 volume ratio matched the ratio of prey mass consumed by the predator species in pilot trials, which we hoped would maximize the chance of detecting prey reduction by both predators (too large a zooplankton:predator ratio and the reduction may not be detected; too small a zooplankton:predator ratio and the predator could quickly deplete the prey and hinder detection of the effect size). We also established control microcosms of both sizes (1.5 L and 100 mL) with no predator. To enable the estimation of overall predator effects over multiple densities (as would be encountered in the field), we halved the zooplankton density in half of all microcosms via filtration. We accomplished this with a 44-μm filter for the 100 mL microcosms, but in the interest of time we used a 100-μm sieve to filter the 1.5 L microcosms. This meant the smallest zooplankton (rotifers) were not reduced in the *Notonecta* microcosms; however, rotifers comprised only 0.46% of zooplankton mass across all control microcosms. All treatment combinations were replicated five times, yielding 40 total microcosms (2 predator species/microcosm sizes × predator presence or absence × 2 zooplankton concentrations × 5 replicates). The microcosms were randomized and placed in an environmental chamber maintained at 25 °C with fluorescent lights on a 16:8 h light:dark cycle. After five days, we filtered the contents of each microcosm using a 44-μm filter and preserved them in 10% Lugol’s solution. We then estimated biomass of the zooplankton taxa as described in Appendix S2.

### Field experiment

To evaluate the influence of predator diversity on phytoplankton biomass and stability, we established replicate pond communities in 200-L cattle tanks at Brackenridge Field Laboratory. Tanks were outfitted with float valves to maintain constant water levels and covered in 1 mm^2^ mesh screens to prevent immigration or emigration of macroorganisms. Before beginning the experiment, we analyzed total N and P of the water following the methods of the American Public Health Association (APHA 1989). We then supplemented NaNO_3_ and NaH_2_PO_4_•H_2_O to bring the total N and P to the concentrations found in COMBO medium (14 mg/L N and 1.55 mg/L P), a nutrient-rich medium commonly used for culturing plankton (Kilham et al. 1998). Immediately following weekly sampling (methods described below) we added both nutrient solutions to compensate for a 5% daily loss rate from the water column (as per Hall et al. 2004), and stirred the water to replenish dissolved oxygen.

We distributed the phytoplankton inoculum equally among the tanks, allowed the phytoplankton to grow for 15 days, and then distributed the zooplankton inoculum equally among the experimental tanks in the same way. Finally, we added to the tanks either no insect predators (controls), 6 adult *Notonecta*, 90 adult *Neoplea*, or half the density of each predator: 3 adult *Notonecta* with 45 adult *Neoplea*. The relative densities of *Notonecta* and *Neoplea* (1:15) were chosen to satisfy the null hypothesis that each tank with predators would experience the same total predation rate if the predators did not partition prey resources. In other words, we used a substitutive design based on consumption rates rather than density *per se*. This predator ratio was derived from the laboratory experiment, where individual *Notonecta* consumed 14.3× and 16.3× more animal mass than *Neoplea* in 0.5× and 1× zooplankton concentration microcosms, respectively. Each treatment was replicated five times, for a total of 20 tanks in a randomized design.

Beginning a week after adding predators, we sampled plankton weekly for six weeks. To sample zooplankton, we used tube samplers to collect ∼6 spatially-spread whole water column subsamples and pool them into a 12-L sample for each tank. We filtered this sample through 65-μm mesh, returned any predators to the tank, and preserved the retained material in 10% Lugol’s solution. To sample phytoplankton, we used 1-cm diameter PVC pipes (one per tank) to collect three spatially-spread whole water column subsamples and pool them into a 50-mL sample for each tank. We estimated biomass of zooplankton taxa as in the laboratory experiment, and additionally estimated biovolume of phytoplankton taxa (methods in Appendix S2). At the end of the experiment, we estimated the remaining density of predators by sampling them with a dipnet until we returned three successive empty sweeps.

### Grouping zooplankton for analysis

To test our prediction that *Notonecta* and *Neoplea* partition zooplankton prey, we divided zooplankton into predicted trophic groups. We expected copepods to be the least vulnerable to predation and therefore placed them in a separate group. Copepods, especially diaptomids (the dominant copepod was the diaptomid *Arctodiaptomus dorsalis*), exhibit a faster escape response and lower vulnerability to predators than other crustacean zooplankton (O’Brien 1979). Copepods are also often omnivorous; in particular, research suggests the other copepod species, *Mesocyclops edax*, is omnivorous as an adult (Adrian and Frost 1993), although diaptomid adults can also be omnivorous (Williamson 1987). We split the remaining zooplankton into two other groups, *Daphnia* and all other smaller zooplankton, based on our prediction that *Notonecta* would selectively prey on *Daphnia* and *Neoplea* would prey on smaller zooplankton.

### Data analysis

To check for differences in predator survival in the field experiment that might confound the predator treatments, we used two-way ANOVA to compare predator survival rates by species, diversity treatment, and their interaction.

We analyzed the effects of the predators on the biomass of zooplankton groups in the laboratory experiment by fitting generalized linear models in the gamma family (gamma GLMs) separately for each predator species and zooplankton group. We first fit the models with the interaction between predator presence and zooplankton concentration (0.5× or 1×) to find whether predator effects were influenced by zooplankton density. Then we fit the models omitting the interaction, with only fixed effects of predator presence and zooplankton concentration so that general predator effects would be assessed across zooplankton densities. Zooplankton groups that were reduced by either predator were then combined as the collective prey for analysis of a diversity effect on zooplankton prey mass in the field experiment. To this end we used the *lme4* package (Bates et al. 2015) to fit a generalized linear mixed model in the gamma family (gamma GLMM) with tank as a random effect to account for repeated measures, and fixed effects for each predator and their interaction. Then we fit all simpler nested models excluding the interaction term, and used likelihood ratio tests to find whether the predator interaction or other terms significantly improved the model fit. If significant, a negative interaction would indicate a synergistic zooplankton reduction while a positive interaction would indicate disruptive effects between the predators. On the other hand, if the interaction did not improve model fit, simple additive predator effects would be supported, if significant. We used the same approach to assess the effect of the predators on the Shannon-Weaver diversity of zooplankton and on mean phytoplankton biovolume. To analyze phytoplankton variability, we calculated the temporal coefficient of variation (CV) of phytoplankton biovolume for each mesocosm. We then repeated the same approach described above to analyze the predator diversity effect, with the exception that no random effect was included in the nested models (i.e., with GLMs rather than GLMMs), as there is only a single CV for each mesocosm. We repeated the phytoplankton analyses excluding *Oocystis*, the largest and likely least edible genus of phytoplankton.

Lastly, we used piecewise structural equation modeling (piecewise SEM) with the *piecewiseSEM* package (Lefcheck 2016) to jointly analyze relationships among all trophic groups. We scaled all variables by their root-mean-square, and used gamma GLMs as the component models. We forced zooplankton to affect phytoplankton CV via its two components, the mean and SD, to lend more insight into the mechanism of any stability effect. Since mean and SD are summary variables (one value per tank), we used time-averaged values of the other variables (zooplankton and predators) to avoid unbalancing sample sizes. We first fit an SEM aligning with the diversity effect analysis above, with collective zooplankton prey combined into one variable and the predator interaction as a predictor of this variable in addition to the individual predators. We then fit a separate SEM with the zooplankton groups separated as in the laboratory analysis, and with no predator interaction. Since adult copepods are considered omnivorous, we first fit the SEM with a pathway from copepods to both the smaller zooplankton and to phytoplankton. To be conservative, we also included *Spirostomum* in the herbivorous smaller zooplankton despite characterization in the literature as a bacterivore, since it may have consumed small phytoplankton. When tests of directed separation were asymmetrical we used the direction with the more conservative *P* value. All analyses were performed in R v. 3.5.3 (R Core Team 2017).

## RESULTS

### Predator populations and plankton community structure

Neither predator reproduced during the field experiment. The median predator survival rate was 79% (SD = 17%), with no significant difference between species (*F*_1,16_ = 0.377, *P* = 0.548), diversity treatment (*F*_1,16_ = 0.225, *P* = 0.642), or their interaction (*F*_1,16_ = 0.003, *P* = 0.956). A large amount of microalgae settled on the bottom of most field tanks, but there was little algal growth on tank walls. The zooplankton composition in the laboratory differed from the average composition in the field; however, the largest difference was in the relative abundance of the copepod *Arctodiaptomus* (median of 10% of zooplankton biomass without predators in the laboratory versus 56% in the field) which was not important for the experiment (see results below). *Daphnia* and the smaller zooplankton comprised a median of 66% and 23% of zooplankton mass in the absence of predators in the laboratory, respectively, versus 27% and 17% in the field, respectively. In both experiments, the smaller zooplankton were dominated by *Scapholeberis kingi*, and all zooplankton species were shared across the experiments except for a few rare rotifers found only in the field experiment (Appendix S1: Table S2). We did not find evidence for any zooplankton colonization events in the field with the potential to affect the results. The most likely colonizers, rotifers, comprised only 0.2% of zooplankton biomass on average in field tanks.

### Effects of predators on plankton groups

In the short-term laboratory experiment, the predators reduced opposing zooplankton groups. *Notonecta* reduced *Daphnia* biomass by 97% averaged over both zooplankton concentrations, without affecting the smaller zooplankton. On the other hand, *Neoplea* reduced smaller zooplankton biomass by 43.0 µg or 52% with the 1× concentration and 46.2 µg or 90% with the 0.5× concentration, without affecting *Daphnia* biomass (Appendix S3 Table S1). Neither predator affected copepod biomass (Fig. 1a). In the field experiment, the predators caused a more than additive reduction of their collective prey (non-copepod zooplankton) when together (Fig. 1b). This is supported by the negative interactive effect of the predators on their collective prey which significantly improved the GLMM (Table 2a, Table 3a). Despite our halving of predator population sizes in tanks with both predators, median zooplankton prey biomass was reduced by 54% with both predators compared to *Notonecta-*only tanks and by 93% compared to *Neoplea*-only tanks. The two predators had additive negative effects on zooplankton Shannon-Weaver diversity (Appendix S3 Table S2, Table S3).

**Table 2.** Results of likelihood ratio tests comparing nested models, including the degrees of freedom (Df), deviance (inverse goodness of fit), and *P* value: **a)** GLMMs for non-copepod zooplankton biomass, **b)** GLMMs for phytoplankton biovolume, and **c)** GLMs for CV of phytoplankton biovolume. *Notonecta* × *Neoplea* is the interaction model with main effects.

| Model | Df | Deviance | *P* |
| --- | --- | --- | --- |
| a) non-copepod zooplankton |  |  |  |
| random intercepts only (null) | 3 | 1263.0 |  |
| *Notonecta* | 4 | 1254.2 | **0.0029** |
| *Neoplea* | 4 | 1262.4 | 1.0000 |
| *Notonecta* + *Neoplea* | 5 | 1253.8 | **0.0033** |
| *Notonecta* × *Neoplea* | 6 | 1248.1 | **0.0168** |
| b) phytoplankton (mean) |  |  |  |
| random intercepts only (null) | 3 | 2877.1 |  |
| *Notonecta* | 4 | 2877.1 | 0.8198 |
| *Neoplea* | 4 | 2877.1 | 1.0000 |
| *Notonecta* + *Neoplea* | 5 | 2877.0 | 0.7697 |
| *Notonecta* × *Neoplea* | 6 | 2875.9 | 0.3004 |
| c) CV of phytoplankton |  |  |  |
| intercept only (null) | 19 | 10.290 |  |
| *Notonecta* | 18 | 9.572 | 0.2238 |
| *Neoplea* | 18 | 9.974 | 1.0000 |
| *Notonecta* + *Neoplea* | 17 | 9.548 | 0.3487 |
| *Notonecta* × *Neoplea* | 16 | 7.662 | **0.0486** |

**Table 3.** Coefficient estimates and standard errors (SE) for models analyzing effects of the predators and their interaction on **a)** non-copepod zooplankton biomass (GLMM), **b)** phytoplankton biovolume (GLMM), and **c)** temporal CV of phytoplankton biovolume (GLM).

| Parameter | Estimate | SE |
| --- | --- | --- |
| a) non-copepod zooplankton |  |  |
| intercept | 7.7513 | 0.5880 |
| *Notonecta* | -0.4006 | 0.1385 |
| *Neoplea* | 0.0005 | 0.0092 |
| *Notonecta* × *Neoplea* | -0.0137 | 0.0053 |
| b) phytoplankton (mean) |  |  |
| intercept | 10.8908 | 0.3917 |
| *Notonecta* | -0.0012 | 0.0923 |
| *Neoplea* | -0.0007 | 0.0061 |
| *Notonecta* × *Neoplea* | 0.0037 | 0.0036 |
| c) CV of phytoplankton |  |  |
| intercept | -0.6181 | 0.3114 |
| *Notonecta* | -0.0301 | 0.0734 |
| *Neoplea* | 0.0033 | 0.0049 |
| *Notonecta* × *Neoplea* | -0.0059 | 0.0028 |

**Figure 1.**
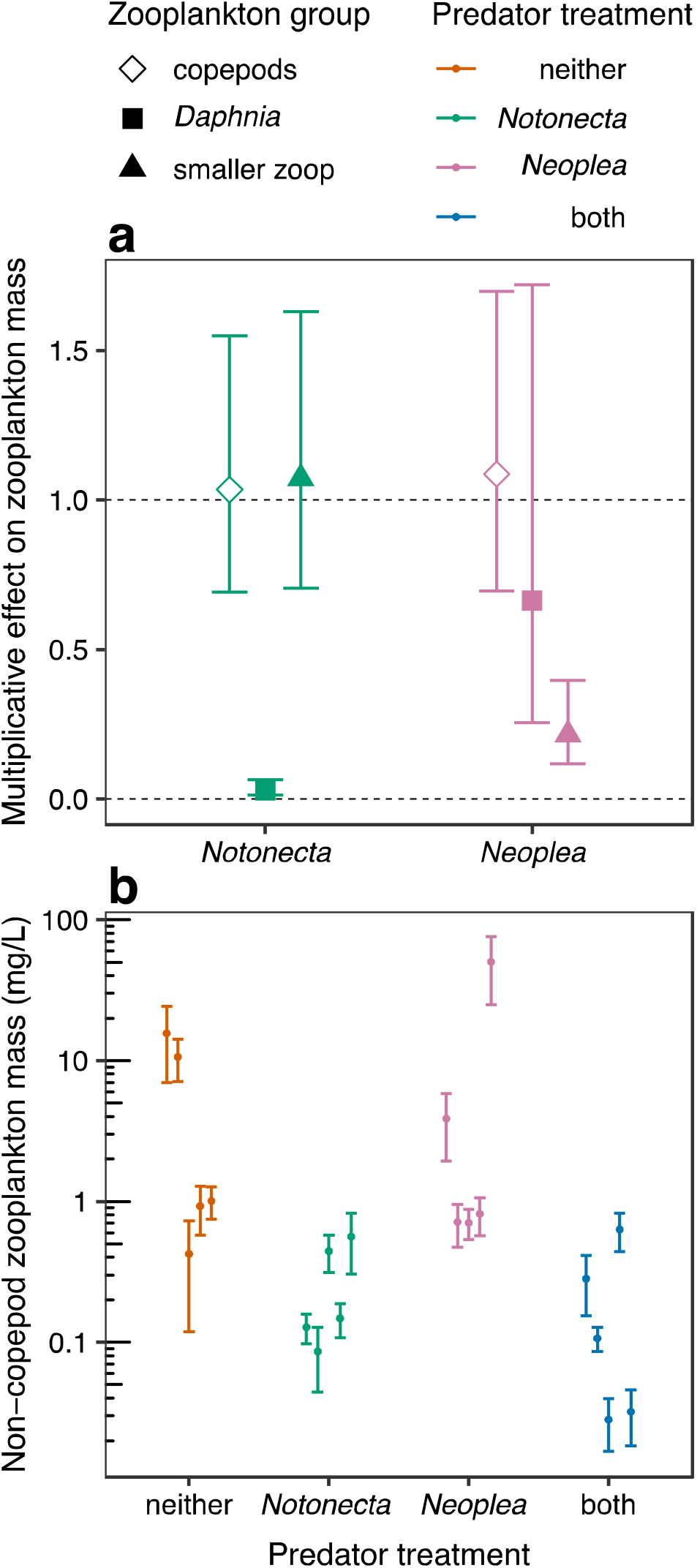
**a)** Multiplicative effects of predator addition on the biomass of zooplankton groups over five days in the laboratory. Shapes are GLM coefficients with 95% confidence intervals. The dashed line at 1 represents no effect, while the dashed line at 0 represents total elimination of a zooplankton group. **b)** Zooplankton dry mass, excluding copepods (summed *Daphnia* and smaller zooplankton), by predator treatment in the field experiment. Dots represent means +/- 1 temporal standard error for each mesocosm. Note the log scale. Also note that predator species densities were halved in the “both” treatment (i.e., substitutive design).

Mean phytoplankton biovolume was not affected by the predators (Fig. 2a, Table 2b, Table 3b). Yet, analogous to the zooplankton prey, the CV of phytoplankton biovolume was synergistically reduced by the predator species (Fig. 2b, Table 2c, Table 3c). Compared to no-predator tanks, phytoplankton CV was 17% lower with *Notonecta*, 34% higher with *Neoplea*, and 52% lower with both predators. Removing the least edible phytoplankter, *Oocystis*, from analysis resulted in a stronger stabilization effect, with phytoplankton CV 20% lower with *Notonecta*, 2% lower with *Neoplea*, and 72% lower with both predators compared to no-predator tanks (Appendix S3 Figure S1, Table S4, Table S5).

**Figure 2.**
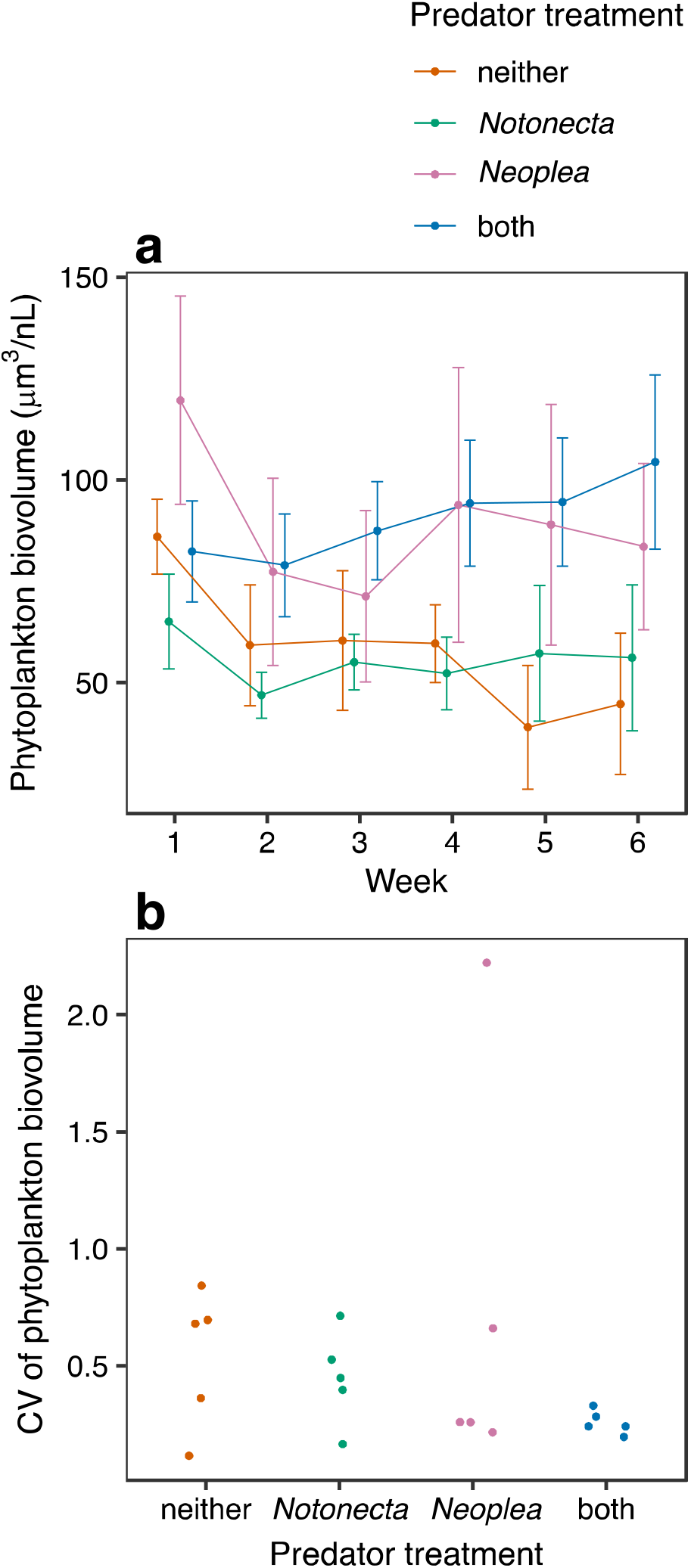
**a)** Time series of mean phytoplankton biovolume, +/- 1 standard error, by predator treatment. **b)** Temporal coefficient of variation (CV) of phytoplankton biovolume by predator treatment (dots represent individual mesocosms).

### SEMs

The piecewise SEM including the predator interaction shows that the cascading diversity-stability effect was mediated by the zooplankton prey and by mean phytoplankton biovolume (Fig. 3). The predators had a negative synergistic (interactive) effect on zooplankton prey, which in turn had a negative effect on the mean, but not the SD, of phytoplankton biovolume. Mean phytoplankton negatively affected phytoplankton CV, as the mean is the denominator of the CV. These three successive negative effects (*Notonecta* × *Neoplea* → non-copepod zooplankton → mean phytoplankton → phytoplankton CV) amount to an indirect negative effect of the predator combination on phytoplankton CV (the product of three negatives is a negative).

**Figure 3.**
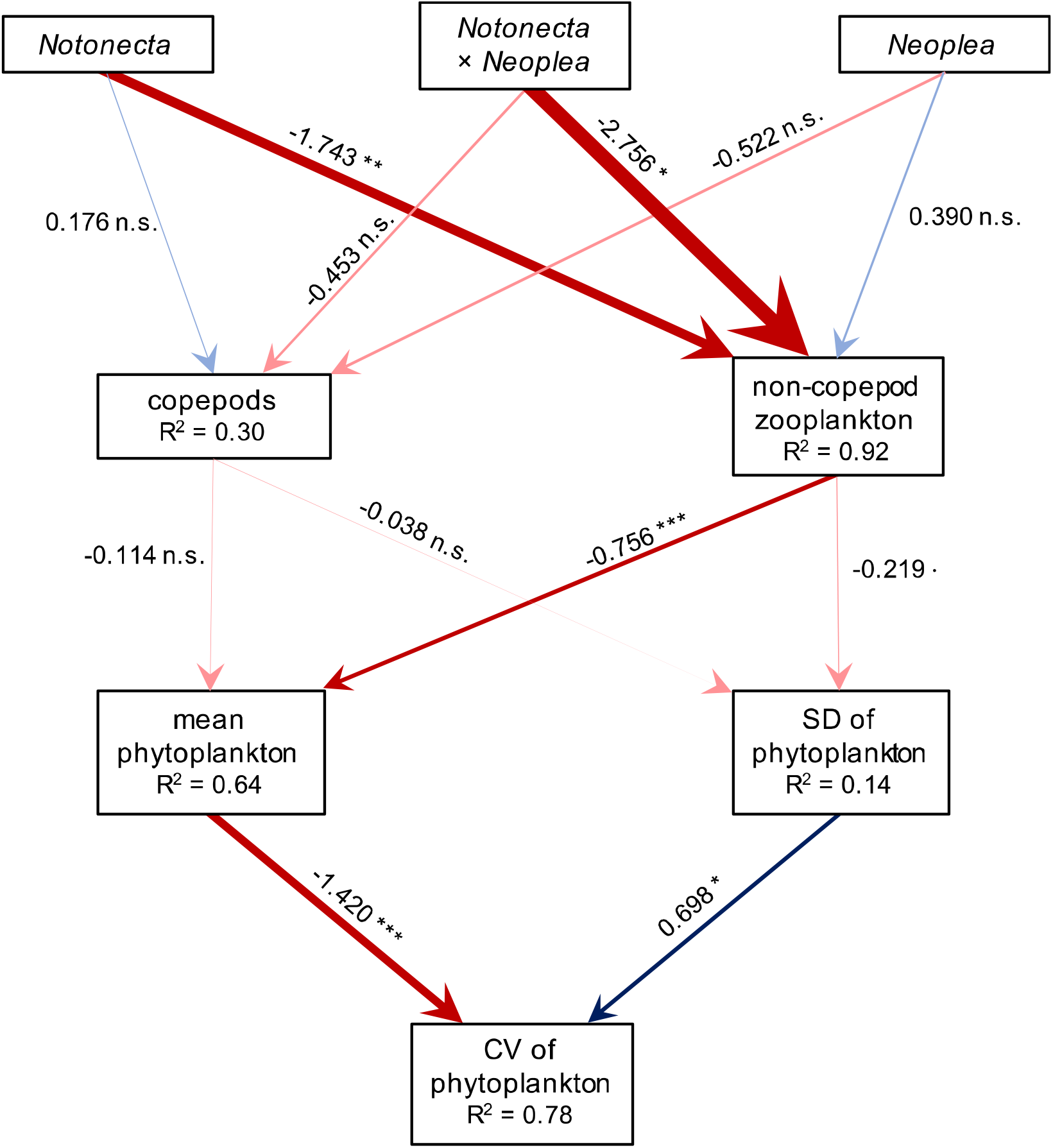
Path diagram for the piecewise SEM with the predator interaction and zooplankton divided into copepods and non-copepods. Red arrows represent negative effects and blue arrows represent positive effects. Arrow thickness is proportional to the coefficient (effect size), which is labeled next to the arrow followed by an indication of the effects’ significance (*** *P*<0.001, ** *P*<0.01, * *P*<0.05, *P*<0.1, n.s. *P*>0.1). The R^2^ value of each component model is provided below its response variable.

With non-copepod zooplankton separated into *Daphnia* and smaller zooplankton, and with copepods considered as omnivores able to consume smaller zooplankton, SEM identified a positive effect of copepods on smaller zooplankton (Appendix S3 Figure S2). Since we did not have a mechanistic explanation for copepods to have a direct positive effect on smaller zooplankton, we concluded that this effect was spurious and was indicative of a positive correlation between the two groups; thus, the pathway from copepods to smaller zooplankton was dropped. The resulting SEM (with copepods as herbivores) suggests that *Notonecta* primarily reduced *Daphnia* while *Neoplea* primarily reduced the smaller zooplankton, although *Notonecta* also reduced the smaller zooplankton to a lesser extent (Fig. 4). Reflecting the non-copepod zooplankton in the previous SEM, it is apparent that *Daphnia* reduced mean phytoplankton. Departing from the previous SEM, however, *Daphnia* also had a weak negative effect on phytoplankton SD, which positively affects phytoplankton CV by definition as its numerator. This negative effect on SD was apparently offset, though, by a slightly stronger positive effect of smaller zooplankton on phytoplankton SD. Thus, the net negative diversity-variability effect as demonstrated in Figure 3 is mirrored in Figure 4 via the pathway *Notonecta* → *Daphnia* → mean phytoplankton → phytoplankton CV. However, the separation of *Daphnia* from smaller zooplankton also revealed a separate pathway contributing to the diversity-stability effect: *Neoplea* → smaller zooplankton → phytoplankton SD → phytoplankton CV, a negative effect followed by two positive effects which amounts to a net negative effect of *Neoplea* on phytoplankton CV.

**Figure 4.**
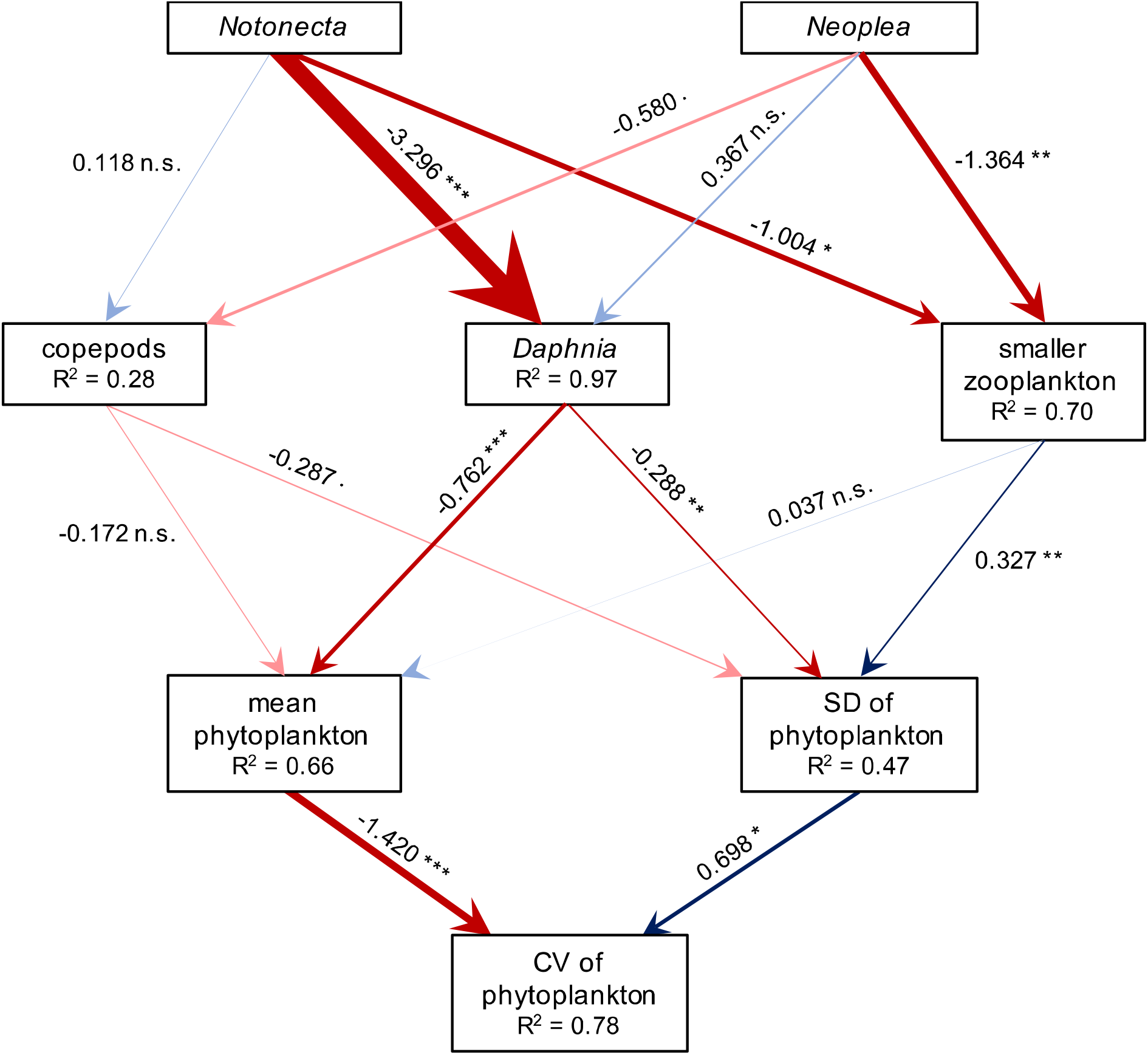
Path diagram for the piecewise SEM with zooplankton divided into three groups: copepods, *Daphnia*, and smaller zooplankton. Red arrows represent negative effects and blue arrows represent positive effects. Arrow thickness is proportional to the coefficient (effect size), which is labeled next to the arrow followed by an indication of the effects’ significance (*** *P*<0.001, ** *P*<0.01, * *P*<0.05, *P*<0.1, n.s. *P*>0.1). Arrows are translucent when *P>*0.05. The R^2^ value of each component model is provided below its response variable.

## DISCUSSION

Our results indicate that the two focal predator species, *Notonecta* and *Neoplea*, partitioned zooplankton prey, leading to a synergistic suppression of zooplankton that stabilized phytoplankton biomass. As revealed by SEM, the stability effect was mediated on the whole by a change in mean phytoplankton biomass without an accompanying proportional change in standard deviation as would normally be predicted by Taylor’s law (Fig. 3; Taylor and Woiwod 1980). Yet, mean phytoplankton biomass was not significantly affected by predator diversity. While these results might seem contradictory on the surface, they can both be correct because SEM lends insight only into likely causal pathways, not into differences among treatments. Given a longer time frame, the predators likely would have reproduced and experienced additional mortality, causing further changes in plankton biomass and stability. However, it is difficult to predict whether such changes would have influenced our conclusions given the dearth of long-term studies integrating biodiversity-ecosystem functioning with food web ecology.

Looking more closely, both predators contributed to the stability effect, despite the stronger overall effect of *Notonecta*. While *Notonecta* lowered phytoplankton CV by nearly eliminating *Daphnia*’s reduction of mean phytoplankton, *Neoplea* lowered phytoplankton CV by suppressing the increase of phytoplankton SD by smaller zooplankton (Fig. 4). Our data cannot answer why *Daphnia* primarily affected mean phytoplankton while smaller zooplankton only affected its temporal variation. However, this difference may be explained by differences in the behavior of these two zooplankton groups. *Scapholeberis kingi*, which comprised 80.4% of smaller zooplankton biomass in field tanks, dwells and feeds almost entirely at the water surface, while *Daphnia* feeds throughout the water column. Arditi and colleagues (1991) demonstrated that this difference causes them to have drastically different effects on phytoplankton, with *Scapholeberis* clearing phytoplankton more slowly than *Daphnia* as the water column remains a refuge from *Scapholeberis* but not from *Daphnia*. While the effect of *Scapholeberis* may not have been strong enough to reduce mean phytoplankton, it may have been enough to create a degree of predator-prey cycling, inducing the phytoplankton to fluctuate as predicted by theory (Volterra 1926, Thébault and Loreau 2005). The predators also both affected zooplankton Shannon-Weaver diversity, but we do not believe this mediated the stability cascade. The predators’ effects on zooplankton diversity were weaker than the effects on zooplankton mass, they were additive rather than synergistic, and they were negative. We would expect such a decline in herbivore diversity to have a de-stabilizing or neutral effect on phytoplankton, not a stabilizing effect (Valone and Balaban-Feld 2019).

Synergistic prey suppression by predator species can be attributed to complementarity or facilitation (Jonsson et al. 2017). Since herbivorous zooplankton compete for phytoplankton resources, predation of only a subset of the herbivores could allow the unaffected herbivores to take advantage of the freed resources and increase their biomass, compensating for the loss of the vulnerable herbivores. In this scenario, if predators individually target a subset of herbivores but the full complement of predators suppresses all herbivores simultaneously, the combination of predators would lead to a synergistic reduction of the herbivores. This scenario aligns with our experiment. It is also possible that the predators facilitated each other, for example if the behavior of one predator disturbed the other predator’s prey from a refuge, thereby increasing its vulnerability to the second predator (Culshaw-Maurer et al. 2020). The lack of refuges in the tanks makes this scenario less likely, however.

The cascading diversity effect was not reflected in all tanks or plankton taxa (i.e., copepods), but the observed pattern was consistent with background variation in the plankton community. In at least one mesocosm per treatment, variability of phytoplankton biomass was at least as low as the average variability with both predators (Fig. 2b). This pattern appears to stem from the large background variation in the system. Even in the absence of predators, variation in the density of *Daphnia* and of the smaller zooplankton was very large, with some predator-free control tanks having consistently low zooplankton densities (Fig. 1b). This meant that only some tanks within each treatment contained enough zooplankton for a cascading predator effect to be detected. Regardless of zooplankton abundance, copepods did not play a part in the cascading effect. Copepods are known to be significantly less vulnerable to predation than other crustacean zooplankton (O’Brien 1979); but why they had little effect on phytoplankton or smaller zooplankton is less clear. It is possible they consumed settled algae, which was not sampled. Similarly, removal of the likely least edible phytoplankton taxon, *Oocystis*, from analysis revealed a stronger stabilization effect on the remaining phytoplankton (Appendix S3 Figure S1, Table S4, Table S5). The fact that less vulnerable taxa played less of a role in the cascading effect adds another line of evidence for the hypothesized consumption-mediated effect.

This study provides a simultaneous evaluation of effects of predator diversity on both the average and variability of primary producer biomass. Most previous predator diversity-ecosystem function experiments measured the mean but not the variability of trophic group biomass as dependent variables (Bruno and O’Connor 2005; Straub et al. 2008). Those experiments indicate that if predators partition their resources, increasing predator diversity can suppress herbivores and lead to higher mean autotroph biomass, i.e., a diversity-biomass trophic cascade (Table 1; Straub et al. 2008). In our field experiment, increasing predator diversity reduced zooplankton prey but did not significantly increase mean phytoplankton biomass. The SEM suggests that only *Daphnia* impacted mean phytoplankton biomass, not the smaller zooplankton (Fig. 4); overall, the effect of the zooplankton on mean phytoplankton biomass was apparently too weak to complete a diversity-biomass trophic cascade. It is not uncommon for herbivores to have weak effects on plant biomass (Maron and Crone 2006). The strength of herbivory is often dampened by the variable food quality of plants, low encounter rates between herbivores and plants, indirect effects, or other factors (Leibold 1989, Borer et al. 2005, Maron and Crone 2006). In this case, the temporal variability of producer biomass was more sensitive to changes in zooplankton biomass, and therefore to predator presence and composition, than was average producer biomass. Future studies will need to explore the generality of this result.

Previous studies have reported the effects of natural enemy diversity on the variability of food web interactions, and some have described theoretical mechanisms relevant to these studies. A few studies used surveys to relate parasitoid richness to the temporal variability of aggregate parasitism rates, finding either no relationship (Rodriguez and Hawkins 2000) or a negative relationship (Tylianakis et al. 2006, Macfadyen et al. 2011). Griffin and Silliman (2011) found that the combination of two predators which exhibited temporal complementarity in attack rates reduced the temporal variability of the total predation rate on a shared prey. Directly relevant theoretical studies are limited, but theory suggests several mechanisms linking diversity and stability in ecosystems generally (Loreau and de Mazancourt 2013). With a two-trophic level model, Thébault and Loreau (2005) showed that decreasing herbivore biomass stabilizes and increases plant biomass. Coupling this finding with the consensus from the literature that complementarity of prey use by predators tends to reduce prey biomass, it follows that predator complementarity may indirectly stabilize (and increase) plant biomass. This food web diversity-stability mechanism could have important implications for ecosystem management and for biological control.

Our results suggest that adding multiple natural enemies to an agro-ecosystem could stabilize fluctuations in crop yields, provided the natural enemies partition pest resources. While a goal of farmers will always clearly be to achieve as high an average yield as possible, achieving consistent yields can be equally important. Thus our study adds another argument for “conservation biological control,” or natural pest control achieved through the conservation of natural enemy biodiversity (Snyder 2019). We showed that diversity of a predator trait known to correlate with prey preference, body size, can be a key factor leading to the cascading stabilizing effect on phytoplankton. Microalgae, especially phytoplankton, are increasingly cultured as a crop for a variety of purposes, from biomass production for biofuels or animal feed, to nutrient removal from wastewaters. However, algal cultivation is not yet widely practiced at the commercial scale, in large part because it has proven too difficult to achieve consistently high algal yields as algae ponds are easily colonized by various herbivorous zooplankton that are difficult or costly to control with mechanical and chemical means (Smith and Crews 2014, Montemezzani et al. 2015). Adding a functionally diverse array of predators to algal cultivation ponds may therefore be a feasible, economical, self-sustaining way to encourage more reliable algal yields. Terrestrial crops also can suffer yield instability associated with multiple pests, and thus may similarly benefit from addition or encouragement of a community of natural enemy species varying in size (Gurr et al. 2012). Diversity of other functional traits related to prey choice may also encourage crop yield stability in a similar fashion and could also be managed in natural enemy communities when relevant information is available, including microhabitat preference, temporal patterns in predation strength, or variance in mode of prey suppression. For example, top-down control can be stabilized by adding or encouraging some predators more active in warm temperatures and some more active in cold temperatures (Griffin and Silliman 2011), or both parasites and predators (Ong and Vandermeer 2015), encouraging consistently low herbivore densities and stable crop yields. While our study emphasizes the importance of complementarity, redundancy is also likely important with increasing temporal and spatial scales to guard against periods of weakened control by, or extirpations of, natural enemies at certain times or locations (Peralta et al. 2014).

Here we have demonstrated that higher predator diversity, and accompanying complementarity of prey use, can cause a chain of effects that cascade down a food web to stabilize the temporal dynamics of basal resource biomass. This connection between predator diversity and primary producer stability is an important step towards joining biodiversity-ecosystem function theory with food web theory, as biodiversity and ecosystem functioning research has only recently begun incorporating energy flows in a food web context (Barnes et al. 2018). Future work is needed to further explore the simultaneous influence of predator diversity, in its various forms, on both the average and stability of community- and ecosystem-level functioning. It is important to uncover whether stability is generally more sensitive to cascading food web effects than is average biomass, or how the two effects are related in different contexts. Predator diversity may also have different effects on other measures of stability, such as resilience. Developing this field at the intersection of biodiversity-ecosystem functioning and food web ecology not only will improve our understanding of the functioning of natural ecosystems and their vulnerability to biodiversity change, but also will provide information that can be used to manage for more reliable crop production and other ecosystem services.

## ACKNOWLEDGEMENTS

Thanks to L. A. Sekula and J. Earwood for help processing samples, to R. Deans for help with insect collection and experiment setup, to S. Duchicela for help with experimental setup, to D. Correa for help with nutrient analysis, and to D. Nobles for providing equipment. Thanks to A. Wolf, R. Decker, D. Cinoglu, S. Ortiz, E. Francis, D. Grobert, and A. Northup for feedback on an earlier version of the manuscript. This research was supported by the Department of Integrative Biology at the University of Texas at Austin and was made possible by the facilities at Brackenridge Field Laboratory.

