## Appendix S1 for "Predator complementarity dampens variability of phytoplankton biomass in a diversity-stability trophic cascade"

**Appendix S1: Relevant information about plankton taxa in this study**

**Table S1.** Phytoplankton morphospecies/taxa found in the field samples and their cell shape and mean cell biovolume. Taxa are listed in descending order of dominance defined by mean biovolume in all samples.

| **Morphospecies/**  **taxon** | **Shape** | **Mean cell biovolume (μm^3^)** | **Notes** |
| --- | --- | --- | --- |
| *Selenastrum* | curved cylinder ending in 2 half-spheres | 22.7 |  |
| *Oocystis* 1 | spheroid | 105 | Effective cell volume = ~1016 μm^3^ (for herbivores) due to mother cell wall |
| *Oocystis* 2 | spheroid | 59.2 | Effective cell volume = ~203 μm^3^ (for herbivores) due to mother cell wall |
| green ovoid 1 | spheroid | 13.9 |  |
| green round | sphere | 12.9 |  |
| green ovoid 2 | spheroid | 21.8 |  |
| *Ankistrodesmus* | cylinder + 2 cones | 2.78 | Often found in clumps consisting of dozens of cells |
| green picoplankton | sphere | 1.83 |  |
| green pennate | rectangular prism ending in two triangular prisms | 27.9 |  |

**Table S2.** Primary zooplankton taxa found in no-predator control samples from the field and laboratory experiments, in descending order of dominance in the field samples. Dominance ranks are determined by median biomass or by mean biomass when the median is 0, and are given for both the field and laboratory experiment for comparison. Eleven taxa not listed here, all rotifers, were rare and represented a negligible contribution to total herbivore mass.

| **Taxon** | **Classification** | **Median mass in field (μg/L)** | **Dominance rank in field** | **Dominance rank in lab** | **Grouping** |
| --- | --- | --- | --- | --- | --- |
| *Arctodiaptomus dorsalis* | Copepoda | 155 | 1 | 3 | copepods |
| *Daphnia magna* | Cladocera | 75.6 | 2 | 1 | *Daphnia* |
| *Scapholeberis kingi* | Cladocera | 34.6 | 3 | 2 | smaller zooplankton |
| *Mesocyclops edax* | Copepoda | 12.9 | 4 | 11 | copepods |
| *Spirostomum* | Ciliophora | 3.88 | 5 | 5 | smaller zooplankton |
| *Euchlanis* | Rotifera | 0.177 | 6 | 13 | smaller zooplankton |
| small ploimid | Rotifera | 0.0632 | 7 | 8 | smaller zooplankton |
| *Monostyla quadridentata* | Rotifera | 0.0375 | 8 | 6 | smaller zooplankton |
| *Monostyla copeis* | Rotifera | 0.0118 | 9 | 9 | smaller zooplankton |
| ostracod | Ostracoda | 0 | 10 | 4 | smaller zooplankton |
| *Chydorus* | Cladocera | 0 | 11 | 7 | smaller zooplankton |
| *Monostyla bulla* | Rotifera | 0 | 12 | 10 | smaller zooplankton |
| larger bdelloid | Rotifera | 0 | 13 | 12 | smaller zooplankton |
