## Appendix S2 for "Predator complementarity dampens variability of phytoplankton biomass in a diversity-stability trophic cascade"

**Appendix S2: Methods for measuring zooplankton biomass and phytoplankton biovolume**

*Zooplankton*

We identified and counted zooplankton in subsamples such that for each taxon, at least 25 individuals or 10% of the sample was counted, whichever came first, and at least 50 total individuals were counted. We measured the length of each crustacean and *Spirostomum* with a micrometer to the nearest half increment (0.24 mm) and measured the length and width of rotifers and width of *Spirostomum* to the nearest 0.05 increment (0.024 mm). We converted crustacean length to dry mass using length-mass regressions (Culver et al. [1985] for copepods, McCauley [1984] for *Scapholeberis*, Dumont et al. [1975] for *Daphnia magna* and *Chydorus*, and Anderson et al. [1998] for ostracods). When rotifers were identifiable to species, we converted rotifer dimensions to dry mass using species-specific equations from the EPA protocol (EPA 2016); otherwise, we used biovolume equations (McCauley 1984) and converted to dry mass assuming a 10:1 biovolume:dry mass ratio. We converted *Spirostomum* dimensions to dry mass approximating cells as cylinders and assuming the same 10:1 biovolume:dry mass ratio.

*Phytoplankton*

We preserved phytoplankton samples in 10% formaldehyde and enumerated phytoplankton densities using a hemocytometer. We counted 25 nL or 50 cells of the most common morphospecies, whichever came first, and counted at least 100 nL for the other taxa. Two individuals separately counted each sample using these methods and then the counts were averaged. We also took several micrographs of each morphospecies across tanks and dates and measured the cell dimensions using ImageJ (Schneider et al. 2012), measuring >15 cells of all but the rarest morphospecies. Then we used geometric approximations to calculate the biovolume of each cell (Appendix S1: Table S1).
