## Appendix S3 for "Predator complementarity dampens variability of phytoplankton biomass in a diversity-stability trophic cascade"

**Appendix S3: Additional results**

**Table S1.** Results of gamma GLMs testing the effect of predator presence, zooplankton concentration (effect of increasing from 0.5× to 1×), and their interaction, for each combination of predator species and zooplankton group in the laboratory experiment.

| **Parameter** | **Estimate** | **SE** | **t** | ***P*** |
| --- | --- | --- | --- | --- |
| **a)** copepods + *Notonecta* |  |  |  |  |
| intercept | 5.8507 | 0.2013 | 29.063 | **<0.001** |
| *Notonecta* | 0.0747 | 0.2847 | 0.262 | 0.797 |
| zoop. concentration | 0.3660 | 0.3020 | 1.212 | 0.246 |
| *Notonecta* * zoop. concentration | -0.0916 | 0.4270 | -0.215 | 0.833 |
| **b)** *Daphnia* + *Notonecta* |  |  |  |  |
| intercept | 7.2364 | 0.3573 | 20.252 | **<0.001** |
| *Notonecta* | -3.2537 | 0.5053 | -6.439 | **<0.001** |
| zoop. concentration | 1.3065 | 0.5360 | 2.438 | **0.029** |
| *Notonecta* * zoop. concentration | -0.6651 | 0.7580 | -0.877 | 0.395 |
| **c)** smaller zoop. + *Notonecta* |  |  |  |  |
| intercept | 7.6501 | 0.2102 | 36.396 | **<0.001** |
| *Notonecta* | 0.1201 | 0.2973 | 0.404 | 0.692 |
| zoop. concentration | 0.2806 | 0.3153 | 0.890 | 0.389 |
| *Notonecta* * zoop. concentration | -0.1147 | 0.4459 | -0.257 | 0.801 |
| **d)** copepods + *Neoplea* |  |  |  |  |
| intercept | 2.9172 | 0.2339 | 12.472 | **<0.001** |
| *Neoplea* | 0.1162 | 0.3308 | 0.351 | 0.730 |
| zoop. concentration | 0.6816 | 0.3308 | 2.061 | 0.056 |
| *Neoplea* * zoop. concentration | -0.0659 | 0.4678 | -0.141 | 0.890 |
| **e)** *Daphnia* + *Neoplea* |  |  |  |  |
| intercept | 5.0849 | 0.4610 | 11.030 | **<0.001** |
| *Neoplea* | -0.1629 | 0.6520 | -0.250 | 0.806 |
| zoop. concentration | 1.2077 | 0.6520 | 1.852 | 0.083 |
| *Neoplea* * zoop. concentration | -0.4957 | 0.9220 | -0.538 | 0.598 |
| **f)** smaller zoop. + *Neoplea* |  |  |  |  |
| intercept | 3.9356 | 0.1949 | 20.193 | **<0.001** |
| *Neoplea* | -2.3356 | 0.2756 | -8.474 | **<0.001** |
| zoop. concentration | 0.4809 | 0.2756 | 1.745 | 0.100 |
| *Neoplea* * zoop. concentration | 1.6028 | 0.3898 | 4.112 | **<0.001** |

**Table S2.** Results of a log likelihood ratio test comparing nested linear mixed effects models for the effect of the predators on zooplankton Shannon-Weaver diversity. Df = degrees of freedom.

| **Model** | **Df** | **Deviance** | ***P*** |
| --- | --- | --- | --- |
| null model (random intercepts only) | 3 | 57.286 |  |
| *Notonecta* only | 4 | 54.584 | 0.1002 |
| *Neoplea* only | 4 | 57.241 | 1.0000 |
| both predators | 5 | 53.285 | **0.0467** |
| both predators + interaction | 6 | 52.828 | 0.4990 |

**Table S3.** Coefficient estimates and standard errors (SE) of the best model in Table S2 above, the linear mixed effects model with effects of both predators on zooplankton Shannon-Weaver diversity but no interaction.

| **Parameter** | **Estimate** | **SE** |
| --- | --- | --- |
| intercept | 0.9039 | 0.1426 |
| *Notonecta* | -0.0621 | 0.0322 |
| *Neoplea* | -0.0023 | 0.0021 |

**Figure S1. a)** Time series of mean biovolume of phytoplankton excluding *Oocystis*, +/- 1 standard error, by predator treatment. **b)** Temporal coefficient of variation (CV) of biovolume of phytoplankton, excluding *Oocystis*, by predator treatment. Dots represent individual mesocosms.

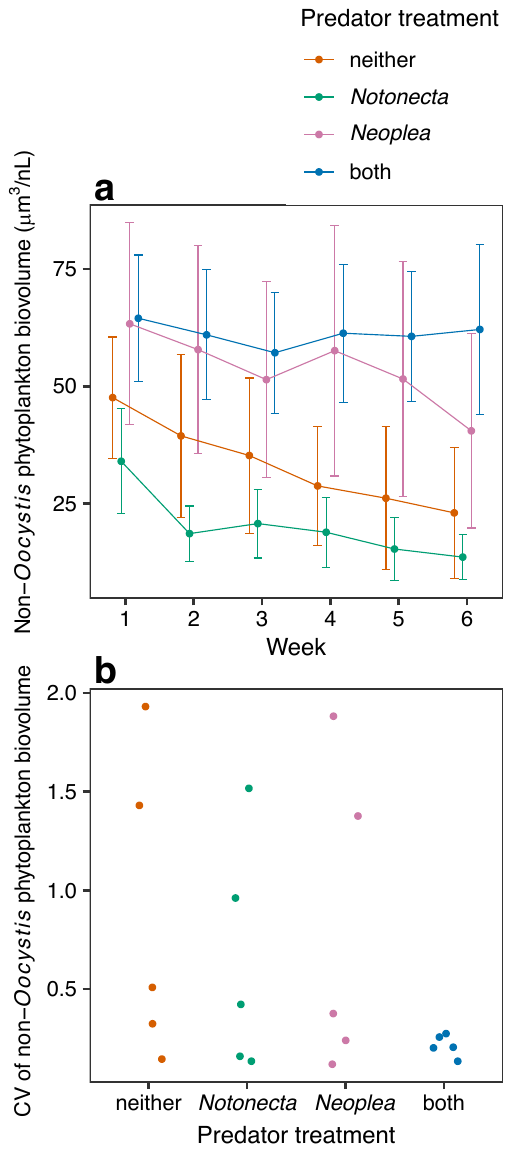

**Table S4.** **a)** Comparison of nested gamma GLMMs evaluating effects of the predators on phytoplankton biovolume excluding *Oocystis*. **b)** Comparison of nested gamma GLMs evaluating effects of the predators on the CV of phytoplankton biovolume excluding *Oocystis*. *Notonecta* × *Neoplea* indicates the full interaction model including both main effects.

| **Model** | **Df** | **Deviance** | ***P*** |
| --- | --- | --- | --- |
| **a)** non-*Oocystis* phytoplankton (mean) |  |  |  |
| random intercepts only (null) | 3 | 2699.7 |  |
| *Notonecta* | 4 | 2699.6 | 0.7492 |
| *Neoplea* | 4 | 2699.7 | 1.0000 |
| *Notonecta* + *Neoplea* | 5 | 2699.6 | 0.7596 |
| *Notonecta* × *Neoplea* | 6 | 2696.9 | 0.1039 |
| **b)** CV of non-*Oocystis* phytoplankton |  |  |  |
| intercept only (null) | 19 | 16.600 |  |
| *Notonecta* | 18 | 16.126 | 0.3752 |
| *Neoplea* | 18 | 16.592 | 1.0000 |
| *Notonecta* + *Neoplea* | 17 | 15.894 | 0.2820 |
| *Notonecta* × *Neoplea* | 16 | 12.168 | **0.0129** |

**Table S5.** Coefficient estimates and standard errors (SE) of the full models in Table S4 above.

| **Parameter** | **Estimate** | **SE** |
| --- | --- | --- |
| **a)** non-*Oocystis* phytoplankton (mean) |  |  |
| intercept | 9.9570 | 0.5727 |
| *Notonecta* | -0.0591 | 0.1350 |
| *Neoplea* | -0.0015 | 0.0090 |
| *Notonecta* × *Neoplea* | 0.0088 | 0.0052 |
| **b)** CV of non-*Oocystis* phytoplankton |  |  |
| intercept | -0.2706 | 0.3473 |
| *Notonecta* | -0.0379 | 0.0819 |
| *Neoplea* | -0.0002 | 0.0055 |
| *Notonecta* × *Neoplea* | -0.0085 | 0.0032 |

**Figure S2.** Path diagram for the piecewise SEM with copepods modeled as omnivores, i.e. able to directly affect smaller zooplankton*.* Red arrows represent negative effects and blue arrows represent positive effects. Arrow thickness is proportional to the coefficient (effect size), which is labeled next to the arrow followed by an indication of the effects’ significance (*** *P*<0.001, ** *P*<0.01, * *P*<0.05, ⋅ *P*<0.1, n.s. *P*>0.1). Arrows are translucent when *P>*0.05. The R^2^ value of each component model is provided below its response variable. Note the significant positive effect of copepods on smaller zooplankton, which we consider an artifact.

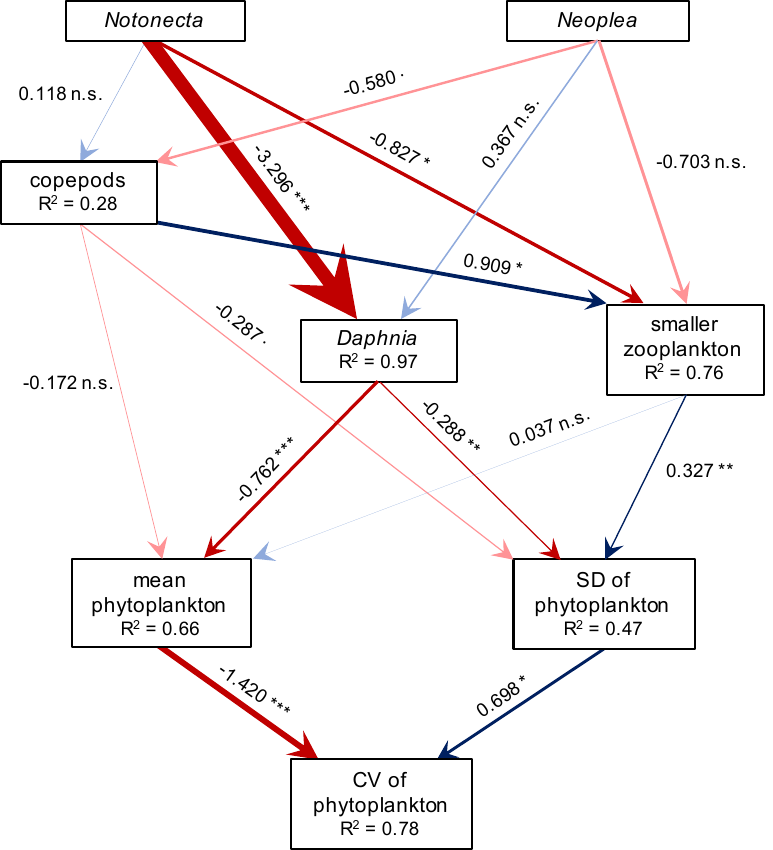
